# Inferring the intrinsic mutational fitness landscape of influenza-like evolving antigens from temporally ordered sequence data

**DOI:** 10.1101/2021.07.28.454153

**Authors:** Julia Doelger, Mehran Kardar, Arup K. Chakraborty

## Abstract

There still are no effective long-term protective vaccines against viruses that continuously evolve under immune pressure such as seasonal influenza, which has caused, and can cause, devastating epidemics in the human population. For finding such a broadly protective immunization strategy it is useful to know how easily the virus can escape via mutation from specific antibody responses. This information is encoded in the fitness landscape of the viral proteins (i.e., knowledge of the viral fitness as a function of sequence). Here we present a computational method to infer the intrinsic mutational fitness landscape of influenza-like evolving antigens from yearly sequence data. We test inference performance with computer-generated sequence data that are based on stochastic simulations mimicking basic features of immune-driven viral evolution. Although the numerically simulated model does create a phylogeny based on the allowed mutations, the inference scheme does not use this information. This provides a contrast to other methods that rely on reconstruction of phylogenetic trees. Our method just needs a sufficient number of samples over multiple years. With our method we are able to infer single-as well as pairwise mutational fitness effects from the simulated sequence time series for short antigenic proteins. Our fitness inference approach may have potential future use for design of immunization protocols by identifying intrinsically vulnerable immune target combinations on antigens that evolve under immune-driven selection. This approach may in the future be applied to influenza and other novel viruses such as SARS-CoV-2, which evolves and, like influenza, might continue to escape the natural and vaccine-mediated immune pressures.

## I. INTRODUCTION

Global seasonal influenza epidemics are caused by influenza A and B viruses that, although being effectively targeted by natural immune responses, seasonal vaccination responses as well as long-term immune memory are able to persistently escape population-wide immunity via mutations [1]. The dominantly targeted antigen of influenza virus is the glycoprotein HA that is located on the viral surface together with the other surface glycoprotein NA, which also acts as antigen. HA is responsible for binding to sialic acid on human cell surfaces and it thereby enables viral cell entry. The human immune system produces antibodies, which primarily bind to different regions (epitopes) on HA thereby blocking the virus from cell attachment and entry. There are 5 dominant and easily accessible epitope regions on the head of HA that have been identified in the circulating subtype H3, which are labeled with the letters A-E [2–4]. These represent the parts of the protein sequence, where the virus predominantly produces amino acid substitutions that abrogate antibody binding and thus lead to immune escape [5].

These interlinked dynamics of the mutating virus and responding human immunity cause a gradual evolution of the viral antigens that is known as antigenic drift [6], which leads to characteristic strain succession patterns in seasonal influenza (Fig. 1). Antigenic drift is also responsible for the fact that there is currently no long-term protective vaccine against seasonal influenza and why still around half a million people die globally from influenza infection [7]. Therefore it is important to create more effective vaccines and other immunization strategies, which target the virus where it is most vulnerable.

**Figure 1.**
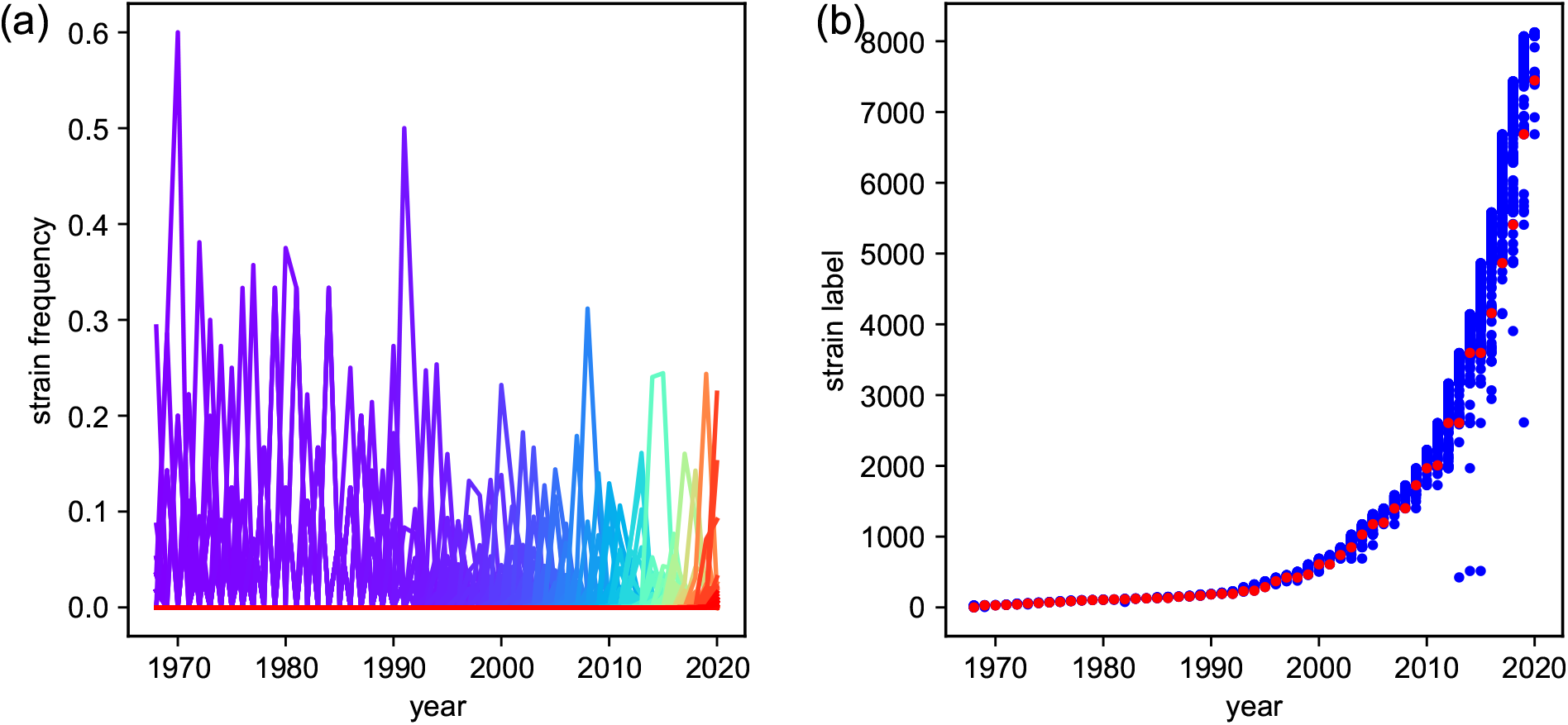
Strain succession for the evolution of HA (H3N2) sequences between 1968 and 2020. (a) Each unique HA amino acid sequence (strain) is shown with its observed frequency in each year as a solid line, with line colors ranging from purple (old strains) to red (new strains). (b) Strains are labeled with increasing numbers from old strains (low labels) to new strains (high labels). The respective strain, which is the most prevalent in each year is marked as red circle. Blue circles indicate strains that were observed with some non-zero frequency.

Even for the currently widely used seasonally updated influenza vaccines, the choice of vaccine strains is not trivial. For best efficacy one needs to make accurate predictions of the viral strains that will be prevalent in the future, based on past and current sequence information. Every year the WHO uses detailed information from international laboratories and worldwide experts to create recommendations on the composition of the influenza virus vaccine [8], but many seasonal vaccines still have a low efficacy compared to other viral vaccines. Thus many computational and experimental efforts are undertaken, which exclusively work on the task of analyzing and predicting the evolution of influenza antigenic sequences, with the goal of making seasonal vaccines more effective [6, 9–14]. But, although periodically updated vaccinations are continually improved and are currently the most effective method for preventive control of seasonal influenza epidemics, such relatively short-term predictions do not generally lead to long-term effective protection of the population [15].

Other approaches aim for cross-protective influenza treatments that are effective against a wide range of strains. Such approaches typically consider strongly conserved epitopes like the receptor binding site (RBS) or the stem of HA [16–25]. Methods targeting those regions require specialized methods for sophisticated vaccine protocols and drug designs [26–36].

Easily accessible sites on highly mutable virus antigens, e.g. on the head of HA in the case of influenza, can generally quickly escape human immune memory via amino acid substitution. However, mutations at some of those strongly targeted sites will be functionally more costly to the virus than others. For a long-term protective immunization approach it therefore would be useful to find and target primarily those easily accessible sites on viral antigens that are most vulnerable, i.e. that have difficulty finding viable mutational escape routes. We can further imagine targeting several sites simultaneously by specifically designed multi-clonal immune responses. In this case it would be useful to choose such combinations of sites as targets, which together are most vulnerable, and do not easily allow the combinations of mutations that lead to escape from the simultaneous responses. The information about the cost of such single and combined mutations at different protein sites is encoded in the intrinsic mutational fitness landscape of the viral sequence.

Previous studies were able to use approaches based on maximum entropy considerations and a method called Adaptive Cluster Expansion (ACE) to computationally infer intrinsic mutational fitness landscapes for other highly mutable viruses, HIV as well as polio, from sequence prevalence data [37–48]. The result of such fitness inference was used to propose a novel cross-protective immunization method against HIV using multidimensionally conserved parts of the proteome, which has been shown to be immunogenic in rhesus macaques [49]. In fact, similar inverse statistical physics models have been extensively used in various context to learn from multiple sequence alignments about the structure and function of various pathogenic and human proteins [50].

Seasonal influenza, however, evolves very differently in the human population than viruses like HIV, for which maximum entropy-based fitness inference methods have been successful. Since influenza is targeted by a population-wide immune memory it is permanently driven away from past strains as opposed to HIV, which evolves much more freely within its fitness landscape and is able to periodically revisit old strains [11, 39, 51]. This immune-driven, non-equilibrium nature of influenza evolution requires a different method for the inference of the intrinsic mutational fitness landscape than the maximum entropy-based methods that were successful for other viruses.

For influenza-like evolving viruses, in which the population-wide immune-memory against each emerging mutant accumulates with every season, the effective fitness landscape depends on the viral evolutionary history and therefore changes in time. Such a changing fitness landscape has also been referred to as a “seascape” [52, 53]. This time-variance of the effective fitness landscape makes approaches that rely on conserved fitness landscapes, such as the recently proposed marginal path likelihood (MPL) method [54], generally unsuitable for fitness inference from sequence time series of influenza-like evolving antigens.

Here we present a method, with which we can infer the single and pairwise mutational intrinsic fitness costs from population-level sequence time series of an influenza-like evolving pathogen. We test our inference approach on sequence data generated by computer simulations and propose its potential application in the future to investigate yearly protein sequence time series data from influenza-like evolving viruses, in order to obtain combinations of vulnerable antibody targets.

## II. MODEL OF INFLUENZA ANTIGEN EVOLUTION

In our model for influenza-like evolution, we consider each epidemic season as an evolutionary step, in which different viral strains, represented as unique protein sequences, evolve and compete with each other according to intrinsic and host immunity-mediated driving forces. In the following we will describe the components of our model, which we use both to create computer-generated sequence data and to motivate the inference method, which we will describe in section III. Our influenza evolution model is motivated and inspired by several previous modeling studies, which describe essential properties of the evolution of influenza-like pathogen populations that lead to the characteristic spindle-like phylogeny and strain succession pattern of seasonal influenza [10, 11, 55–57]. Those pathogen models also relate to more general models of rapid adaptation in asexual populations that evolve towards increasing fitness in a traveling-wave type manner [58–60].

### A. Sequence representation

For the representation of viral strains we use a binary sequence representation, in which a strain 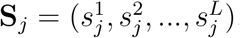, i.e. a unique sequence, is represented as a string of *L* ones and zeros. This is a coarse-grained representation of a real protein, wherein, in principle, there could be 20 possible amino acids at each residue. For proteins that do not mutate too much (like the p24 structural protein of HIV), a binary Ising like representation, instead of a Potts model, is reasonable [40]. Also, our approach could be generalized to Potts models. Here, we consider sequences of *L <* 100, which is much shorter than real protein sequences.

### B. Fitness model

The time-dependent fitness landscape in our model, which defines the fitness of different strains, is composed of two components. The intrinsic fitness represents the intrinsic abilities of viruses belonging to a given strain to infect, reproduce and transmit in a susceptible human population. The host immunity-mediated fitness cost, on the other hand, represents the accumulated immunity against viruses belonging to a given strain in the host population, which reduces the number of susceptible hosts and therefore reduces the fitness of the respective strain. The total fitness of a strain **S**_*j*_ at time *t*, given the evolutionary history **x**(*t*^*′*^ *< t*) of the whole virus population in humans, is modeled as:

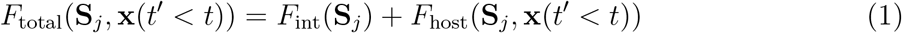

with intrinsic and immunity-mediated fitness components *F*_int_ and *F*_host_.

#### Intrinsic fitness model

The intrinsic fitness of a strain in our model is represented with a 2-point approximation as

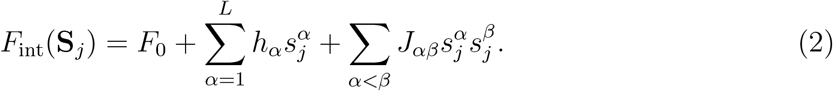

Here *F*_0_ represents the intrinsic fitness of a reference strain which is represented as a string of zeros, the second term represents the fitness change due to independent mutations at each sequence site *α* compared to the reference strain (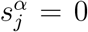 if unmutated, 1 otherwise), and the last term represents the additional fitness change due to coupled mutations at pairs of sites *α* and *β*. The single-mutational fitness coefficients {*h*} and the mutational coupling coefficients {*J*} describe the intrinsic mutational fitness landscape, which we ultimately want to infer from the observed sequences. The intrinsic fitness coefficients describe how easy or difficult it is for the virus to create escape mutations if specific sites or pairs of sites are targeted by the host. Note, that by using this Ising-type approximation of the intrinsic fitness landscape, we reduce the number of fitness parameters for binary sequences with e.g. length *L* = 20 from 2^*L*^ = 1048576 unique strains to *L* * (*L* + 1)*/*2 = 210 fitness parameters {*h, J*}. The fitness model used in Eq. (2) is different compared to maximum entropy models wherein the fitness is the exponential of an expression like Eq. (2). We use this formulation for convenience and for demonstrating our method.

#### Representation of host immunity-mediated fitness costs

The host-dependent immunity-mediated fitness component depends on the evolutionary history of the viral population and in our model is calculated with a functional form similar to that previously used in other influenza fitness models [10, 55, 57], i.e.

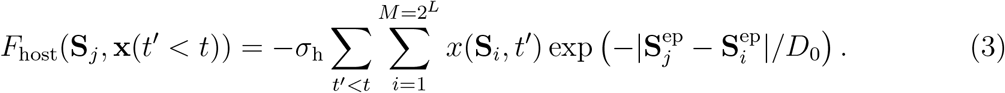

The immunity-mediated fitness component decreases the fitness of each emerging strain over time and is proportional to the prevalence *x*(**S**_*i*_, *t*^*′*^) of antigenically similar strains **S**_*i*_ in previous years *t*^*′*^. This accumulating fitness cost forces the virus to continuously evolve away from previously prevalent sequences. Here 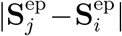 describes the mutational distance between strain **S**_*i*_ and **S**_*j*_ within their immune-targeted epitope regions and *D*_0_ is the crossimmunity distance (i.e., the typical mutational distance within epitope regions beyond which two strains are dissimilar enough to not be targeted by immune responses that were raised against the other). In the following we will assume for simplicity that all modeled sequence sites are equally immune-targeted and therefore 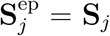, but the model can, in principle, be extended to account for less or untargeted sites in the model sequence **S**_*j*_.

### C. Sequence selection

During the spread of viral infections in the course of a flu season, different strains are assumed to grow with a growth rate given by their respective fitness (Eq. 1), i.e.,

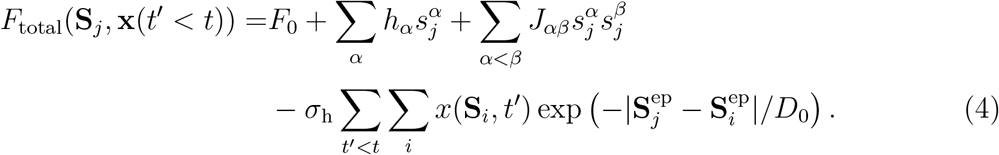

At the end of a season a fixed number *N*_pop_ of sequences is assumed to survive into the next season. The expected frequency of a given strain **S**_*j*_ among the selected sequences in season *t* + 1 is calculated as

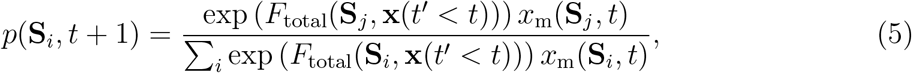

where *x*_m_(**S**_*j*_, *t*) denotes the frequency of strain **S**_*j*_ in season *t* before growth and selection. The number of selected sequences *N* (**S**_*j*_, *t* + 1) belonging to strain **S**_*j*_ are drawn from a multinomial distribution with probabilities given by Eq. (5) and *N*_pop_ as the number of draws.

### D. Sequence mutation

We assume that mutation is a separate step from selection in each flu season. Thus in every modeled season *t* before growth and selection, sequences are modeled to mutate and thereby create a new frequency distribution **x**_m_(*t*). We assume one symmetric mutation rate *µ*, per season, between the two different states at each site, such that the mutation probability *µ*_*ij*_ = *µ*_*ji*_ for mutation between strains **S**_*i*_ and **S**_*j*_ is given as

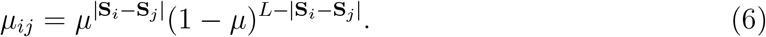

In a stochastic simulation procedure, the mutated sequences can simply be created by randomly switching the state at each site in each selected sequence with probability *µ*.

As mentioned before, the main motivation for the model is to generate a controlled dataset that can be used to develop and test a method for inferring the intrinsic mutational fitness landscape of influenza-like evolving antigenic sequences. The goal within our model framework is to infer the intrinsic fitness coefficients {*h, J*} from yearly observations **x**(*t < T*) (until the most recent season *T*) of antigenic protein sequences, in order to learn about the vulnerability and mutational escape likelihood at different single and combinations of sequence sites upon being targeted.

On this account we developed an inference approach, which we test on computer-generated data that we produced via simulation of our sequence evolution model with a known fitness landscape.

## III. ANALYSIS AND INFERENCE BASED ON SIMULATED SEQUENCE DATA

### A. Simulation produces influenza-like antigen evolution

Based on the presented model we ran stochastic simulations to compare the computer-generated sequence evolution to influenza sequence data and to test our fitness inference method. The simulation parameters are the sequence length *L*, population size *N*_pop_, number of simulated seasons *N*_simu_, mutation rate *µ*, cross-immunity distance *D*_0_, host-immunity coefficient *σ*_h_ and intrinsic fitness coefficients {*h, J*} (cf. Tab. I). In the beginning of each simulation the population is initialized with the unmutated strain **S**_0_ = (0, 0, …, 0). Accordingly the initial strain frequency distribution is given by *x*(**S**_0_, *t* = 0) = 1. The intrinsic fitness landscape in our simulations is predetermined by the chosen intrinsic fitness parameters {*h, J*}. As just an example, we sample a limited number of these parameters from Ising coefficients inferred for the HIV p24 protein using a maximum entropy model [40]. In each time step representing one epidemic season, sequences are first mutated according to rate *µ*. After mutation, current fitness of each present strain is calculated with Eq. (4), based on which the selection probability (Eq. (5)) of each strain is determined. The sequence population for the next season is then sampled by *N*_pop_ random draws from a multinomial distribution with the individual probabilities given by the respective selection probabilities of each strain.

For a range of parameter choices our stochastic simulations produce immune-driven evolutionary patterns (Fig. 2), which are qualitatively similar to those observed for evolution of the influenza spike protein HA (H3N2) in the human population (Fig. 1). This similarity indicates that our model is able to capture the essential dynamics of antigenic evolution for pathogens like seasonal influenza. One difference in the shown figures (Figs. 1b and 2b) is the approximately exponential increase of total sequence diversity in strains based on full HA amino acid sequence data versus the more linear increase of total sequence diversity in a simulation of binary sequences of length 20. The dependence of this growth of strain diversity on various parameters and its underlying mechanisms should be further investigated when translating our procedures to infer the fitness landscape of influenza. We speculate that the exponential increase of sequence diversity in observed influenza sequences may be due to the rapid increase in the amount of yearly acquired sequencing data in the past years.

**Figure 2.**
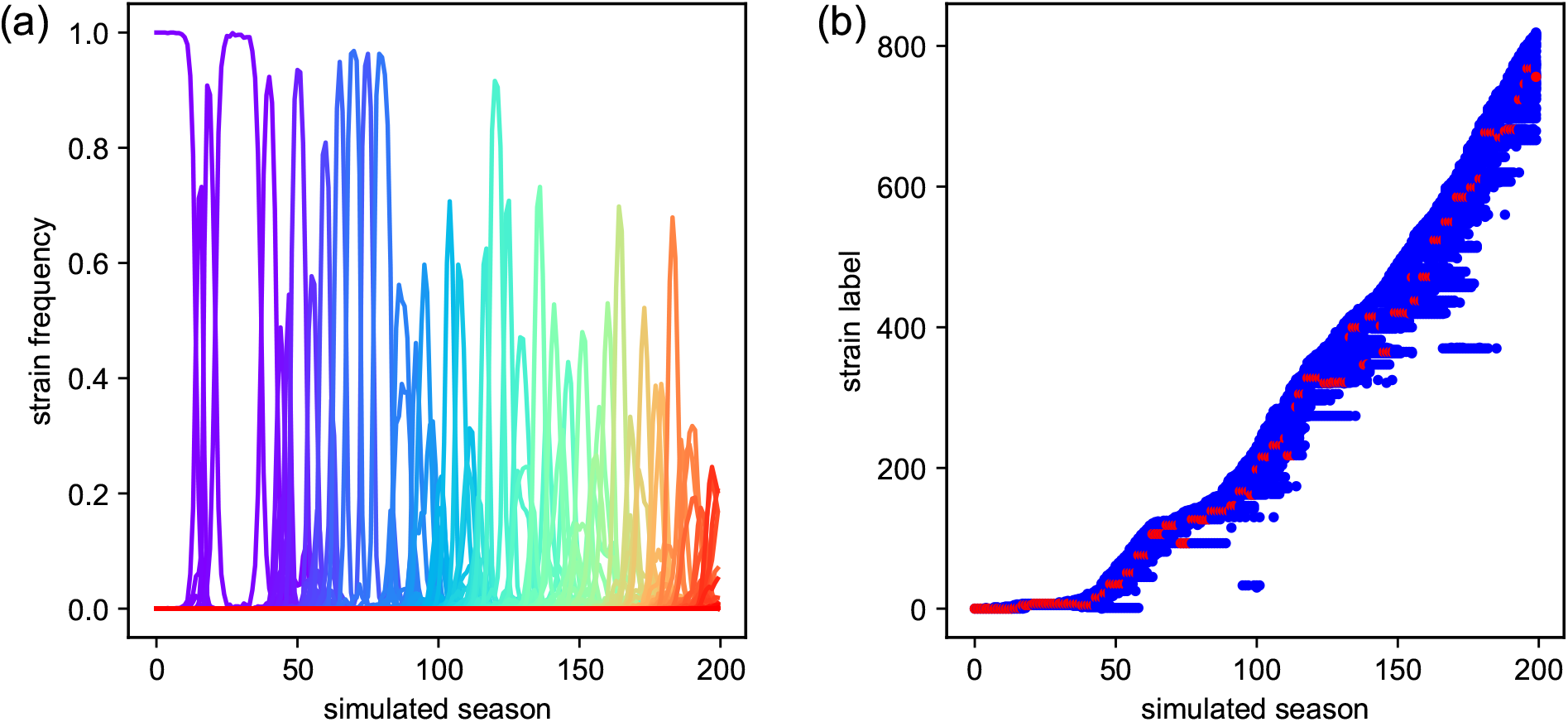
Strain succession for the evolution of simulated data over 200 time steps. (a) Each unique sequence (strain) is shown with its observed frequency in each simulated season as a solid line, with line colors ranging from purple (old strains) to red (new strains). (b) Strains are labeled with increasing numbers from old strains (low labels) to new strains (high labels). The respective strain, which is the most prevalent in each simulated season is marked as red circle. Blue circles indicate strains that were observed with some non-zero frequency. For the shown example the parameter values for simulation and analysis are: *N*_pop_ = 10^5^, *L* = 20, *µ* = 10^*−*4^, *σ*_h_ = 1, *D*_0_ = 5, *N*_simu_ = 200, *B* = 10^3^.

### B. Observation of stringent selection regime

For the analysis of the simulated sequences we randomly sampled a number *B* of sequences per season to imitate the sampling properties of real observed protein data, which contain only subsets of the yearly circulating viruses. For an example set of sampled data from one simulation we see that the distribution of total fitness is more narrow than the distributions of the intrinsic and the immunity-dependent fitness components (Fig. 3). The narrow total fitness distribution in each season indicates a stringent selection regime, in which only those strains in a narrow fitness range around the currently fittest strain survive into the next season. In this observed regime we have

**Figure 3.**
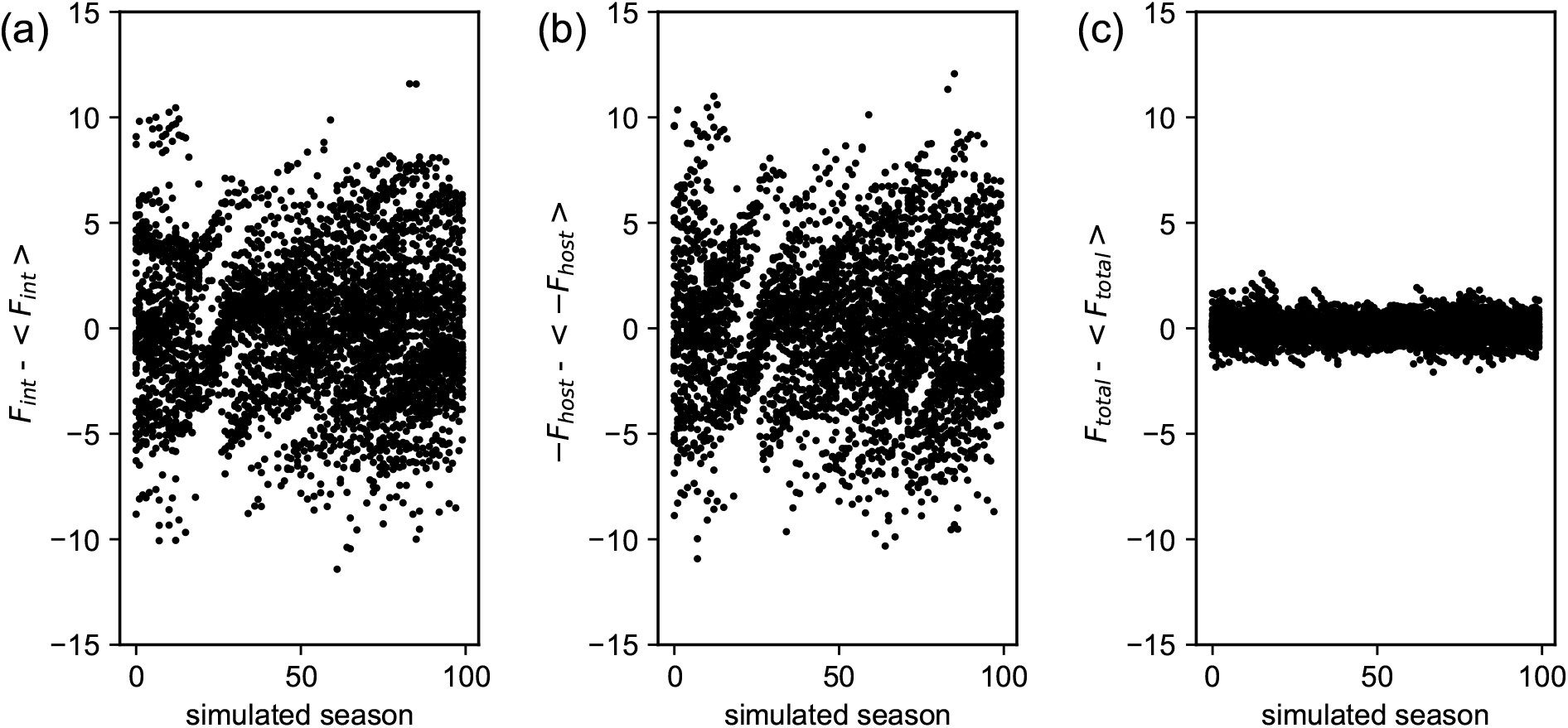
Fitness deviations from the mean of sampled strains for each simulated season between season 100 and 200. (a) Intrinsic fitness component *F*_int_, (b) immunity-dependent fitness component *F*_host_, (c) total fitness *F*_total_ = *F*_int_ + *F*_host_. For the shown example the parameter values for simulation and analysis are: *N*_pop_ = 10^5^, *L* = 20, *µ* = 10^*−*4^, *σ*_h_ = 1, *D*_0_ = 5, *N*_simu_ = 200, *B* = 10^3^.

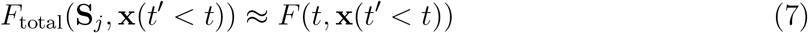

with *F* (*t*, **x**(*t*^*′*^ *< t*)) being a constant for each season *t*, conditional on the specific evolutionary history **x**(*t*^*′*^ *< t*).

Indeed we find with our simulation a clear 1:1 correspondence between the intrinsic fitness variation and immunity-dependent fitness variation in each season (Fig. 4a), which add up to a roughly constant total fitness in each season, as the stringency assumption (Eq. (7)) suggests. In Fig. 4b it can be observed that in our simulation the absolute population fitness decreases with each year, both due to the emergence of less intrinsically fit strains and due to the population-wide accumulation of immune pressure.

**Figure 4.**
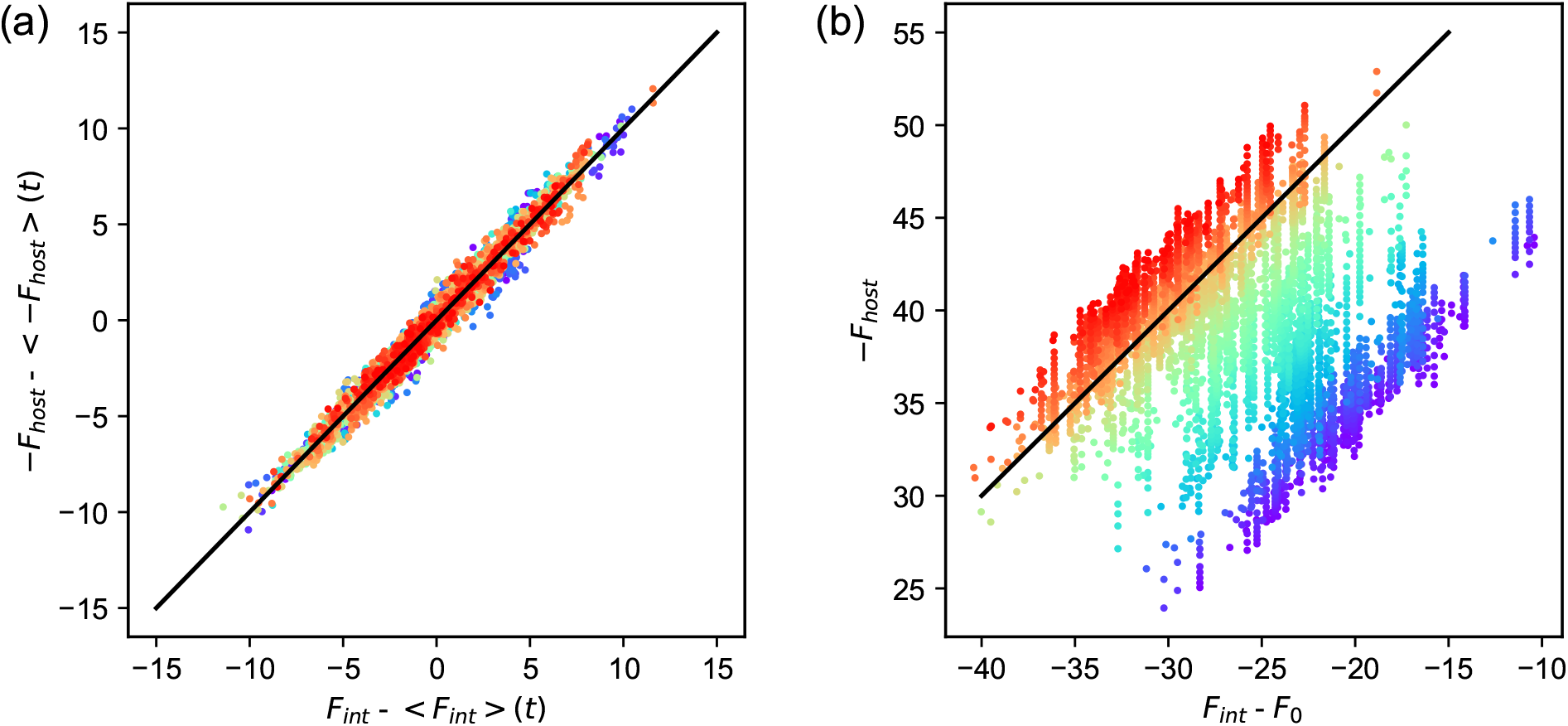
Negative immunity-dependent fitness cost −*F*_host_ (y-axes) compared against intrinsic fitness *F*_int_ (x-axes) for sampled strains for each simulated season between season 100 and 200 (same data as in Fig. 3). (a) Fitness deviations from the mean as colored circles with a solid black line indicating 1:1 correspondence. (b) Absolute fitness components as colored circles with a solid black line indicating slope 1. Colors from purple to red in both panels indicate seasons from 100 to 200, in which the respective strains were sampled.

As for the evolution of influenza in the human population, its seasonal dynamics has been well described with traveling wave models, which indicate a localized, narrow distribution of the viral population in fitness space at any given time point [58–61]. Such a narrow fitness distribution of concurrently selected viral antigenic sequences indicates that one necessary condition (Eq. (7)) for our stringency-based inference method may be fulfilled by seasonal influenza antigens.

### C. Method for intrinsic fitness inference

From Eq. (1) together with Eq. (7) we obtain the following relation for the observed strains **S**_*j*_ in each given year in the case of stringent selection, i.e.,

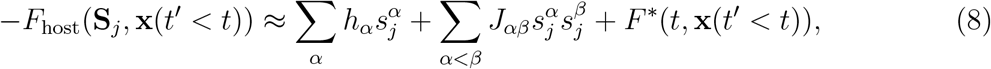

where *F* ^*\**^(*t*, **x**(*t*^*′*^ *< t*)) = *F*_0_ − *F* (*t*, **x**(*t*^*′*^ *< t*)) is another constant at time *t*, conditional on the evolutionary history until *t*. If we approximate the evolutionary history **x**(*t*^*′*^ *< t*) with the observed strain frequencies starting from the first year of observation and assume the model parameters *σ*_h_ and *D*_0_ to be known, e.g. as fit parameters to independent cross-immunity studies [10], we can calculate *F*_host_(**S**_*j*_, **x**(*t*^*′*^ *< t*)) for each observed strain in each season. We now use these host-dependent fitness values together with Eq. (8) to infer the intrinsic fitness coefficients {*h, J*} as well as the additional parameters {*F* ^*\**^} (one additional parameter per season). We here treat {*F* ^*\**^} as independent parameters, although they generally depend on other model parameters and on the history via the full evolutionary dynamics of the system. For the regression we minimize the sum of squared residuals between the data *Y*_data_(**S**_*j*_, *t*) given by the LHS of Eq. (8) and the model *Y*_model_(**S**_*j*_, *t*, {*h, J, F* ^*\**^}) given by the RHS of Eq. (8), i.e.,

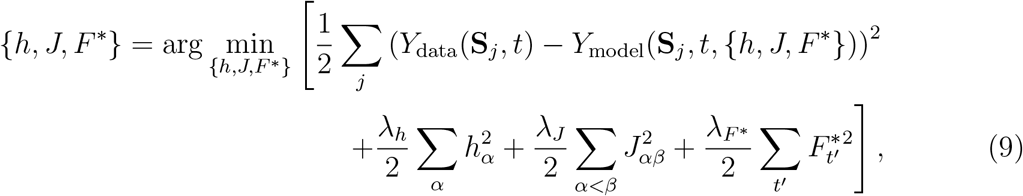

where we also take into account regularization with coefficients *λ*_*h*_, *λ*_*J*_, *λ*_*F\**_ that in a Bayesian sense correspond to Gaussian prior distributions.

For inference we use the following equation, [62, Eq. (3.44)],

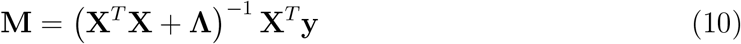

to solve for the unique parameter values 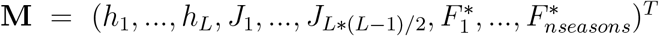, which minimize the sum of squared residuals subject to ridge-regularization (Eq. (9)). The feature vector for each sampled strain, which forms a row in the feature matrix **X**, consists of binary features representing the single-mutational and double-mutational states of the respective sequence as well as its time of observation. **y** is a column vector, whose entries are given by the values −*F*_host_(**S**_*j*_|**x**(*t*^*′*^ *< t*)) (cf. Eq. (3)) for the respective sequence **S**_*j*_, sampled at time *t*. The non-zero regularization coefficients, {*λ*_*h*_, *λ*_*J*_ , *λ*_*F \**_}, are collected in the diagonal matrix **Λ**, and regularization also ensures that no singularities are encountered at matrix inversion. The coefficients *λ*_*h*_ and *λ*_*F \**_ are set to very small values corresponding to a very wide, rather non-restrictive, prior distribution, while *λ*_*J*_ , corresponding to the assumed sparse mutational couplings, is set to 1.

### D. Inferring the intrinsic mutational fitness landscape from simulated influenza-like sequence data

The parameters for simulation and inference with chosen default values are collected in Tab. (I).

**Table I.** Parameters for simulation of influenza-like sequence evolution and for intrinsic fitness inference

| Parameter | Description | Default value |
| --- | --- | --- |
| $\{h, J\}$ | intrinsic fitness coefficients for single mutations and pairwise mutational couplings | sampled values from HIV protein p24 |
| $L$ | length of sequence representation | 20 |
| $\mu$ | mutation rate (per sequence site) | $10^{-4}$ |
| $D_0$ | cross-immunity distance | 5 |
| $N_{\text{pop}}$ | population size | $10^5$ |
| $\sigma_h$ | host-fitness coefficient | 1 |
| $\{\lambda_h, \lambda_J, \lambda_{F^*}\}$ | regularization coefficients for inference | $\{10^{-4}, 1, 10^{-4}\}$ |
| $n_{\text{seasons}}$ | number of years/seasons used for inference | 100 |
| $B$ | number of sampled sequences per year | $10^3$ |

In Fig. 5 we compare the inferred and the simulated intrinsic fitness coefficients for one simulation. The correlation coefficients between simulated and inferred coefficients and in particular the Pearson correlation *r*_*hJ*_ between the total fitness effects of double mutations indicates if the specific fitness inference on the particular sequence data set can successfully distinguish between pairs of sites, at which escape mutations lead to either low or high (negative) fitness costs.

**Figure 5.**
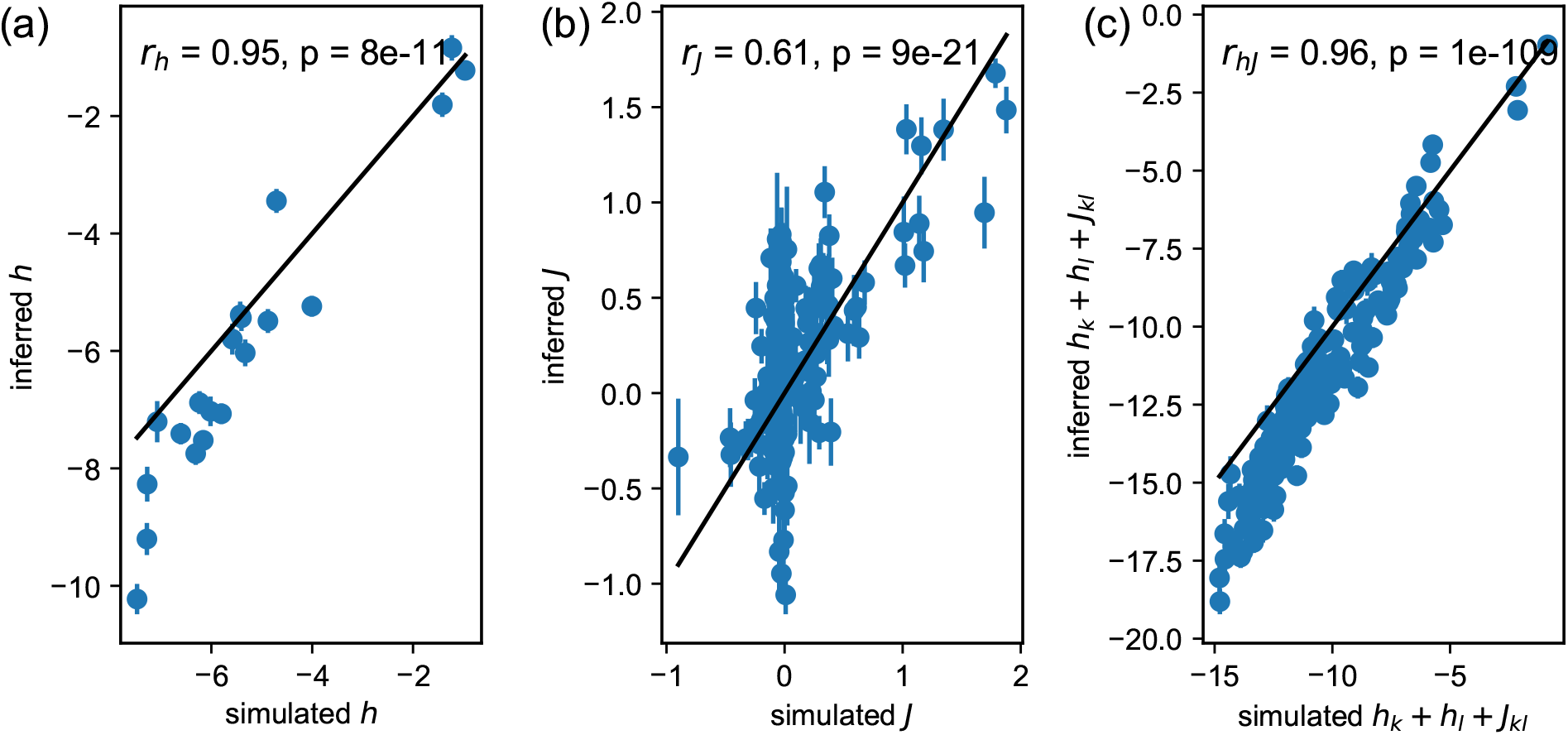
Parameter correlations for the inference on one simulated data set. Inferred values of the fitness coefficients are shown against the fitness coefficients that were used as input values for the simulation. (a) single-site mutational fitness coefficients *h*, (b) coupling coefficients *J* for simultaneous mutations at any two sites, (c) total fitness changes *h*_*k*_ + *h*_*l*_ + *J*_*kl*_ due to simultaneous mutations at any two sites *k* and *l*. Pearson correlation coefficients *r* together with their respective p values are shown in each panel for the respective set of parameters. For the shown example the parameter values for simulation and analysis are: *N*_pop_ = 10^5^, *L* = 20, *µ* = 10^*−*4^, *σ*_h_ = 1, *D*_0_ = 5, *N*_simu_ = 200, *B* = 10^3^, *n*_seasons_ = 100, *λ*_*h*_ = 10^*−*4^, *λ*_*J*_ = 1, *λ*_*F*_ *\** = 10^*−*4^.

Besides the correlation coefficient *r*_*hJ*_ we use another measure for inference performance, which can be useful if we are mainly interested in identifying those pairs of sites that have the most deleterious fitness effect, i.e. those whose intrinsic fitness change compared to the reference sequence is below a certain negative threshold with

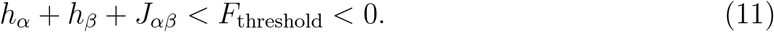

In this case we can use typical classification performance measures to assess how well our inference method can distinguish between deleterious and more neutral or beneficial double mutations. We compare the classification of each pair (based on the inferred coefficients) with the classification of the simulation input values by calculating the precision-recall curve (PRC) as well as the receiver operating characteristic curve (ROC) and the respective areas under the curves (AUC) (Fig. 6), which approach 1 in the case of perfect classification skill.

**Figure 6.**
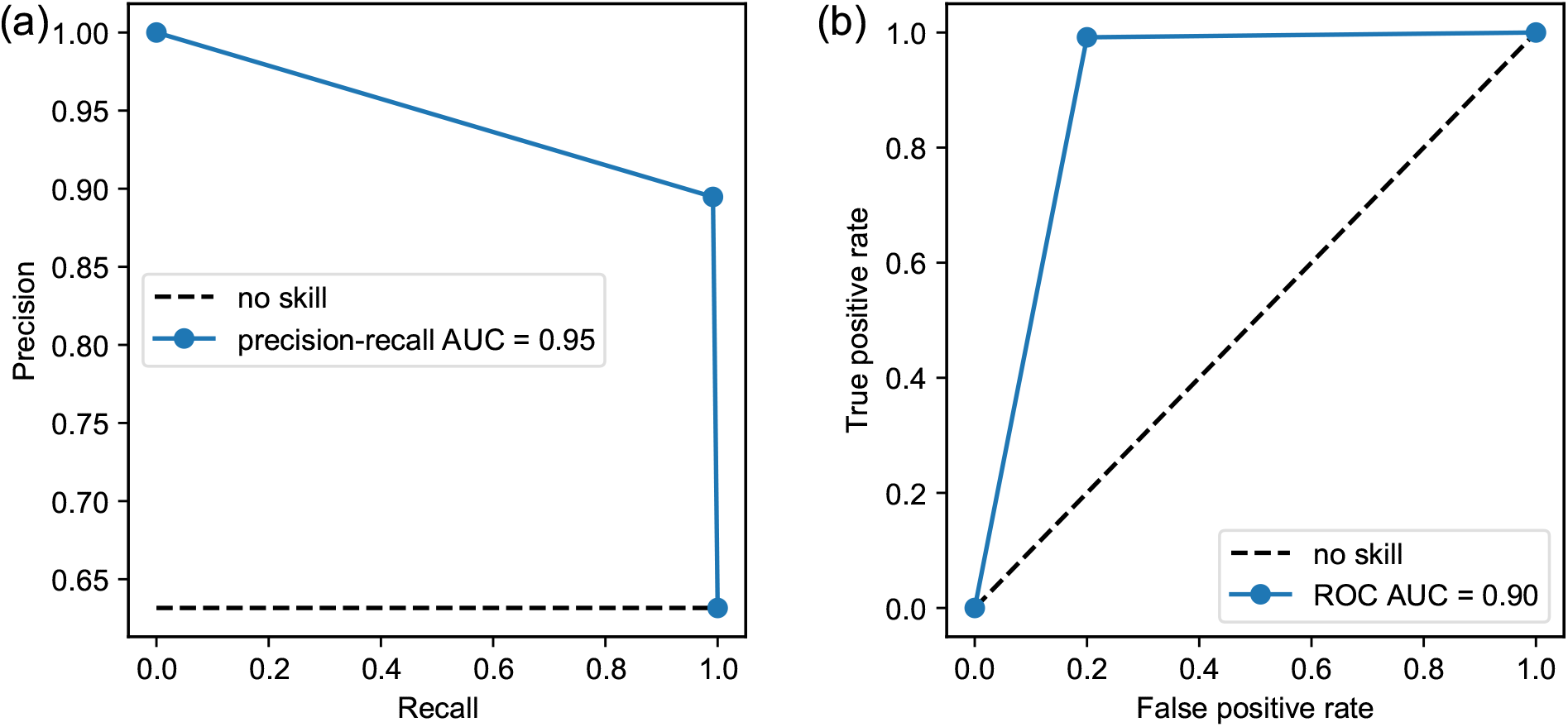
Classification performance for the inference on one simulated data set. Double mutations are classified as deleterious if their total fitness cost is lower than *F*_threshold_ = −10 (cf. Eq. (11)). The precision-recall curve (PRC) and (b) the ROC curve for the classifier derived from inferred fitness coefficients. Black, dashed lines show a no-skill classifier for comparison and the area under the classifier curve (AUC) is given in each panel, respectively. For the shown example the parameter values for simulation and analysis are: *N*_pop_ = 10^5^, *L* = 20, *µ* = 10^*−*4^, *σ*_h_ = 1, *D*_0_ = 5, *N*_simu_ = 200, *B* = 10^3^, *n*_seasons_ = 100, *λ*_*h*_ = 10^*−*4^, *λ*_*J*_ = 1, *λ*_*F*_ *\** = 10^*−*4^, *F*_threshold_ = −10.

When calculating the inference performance for one simulation with sequence length *L* = 20 in terms of correlation *r*_*hJ*_ and classification performance (AUC) for various sample sizes (Fig. 7), we find that a minimum total number of sampled strains *n*_seasons_ \**B* is required for accurate inference. In the shown example a total sample size of ≥ 10^5^ strains is required for high inference performance (Fig. 7b). Since this is true for a sequence of length *L* = 20, a very large number of sequences would be needed for an inference based on a protein representation with all amino acid sites *L >* 100, which indicates that for real proteins sequence representations with strongly reduced dimensions are needed for inferences based on the available amount of observed data.

**Figure 7.**
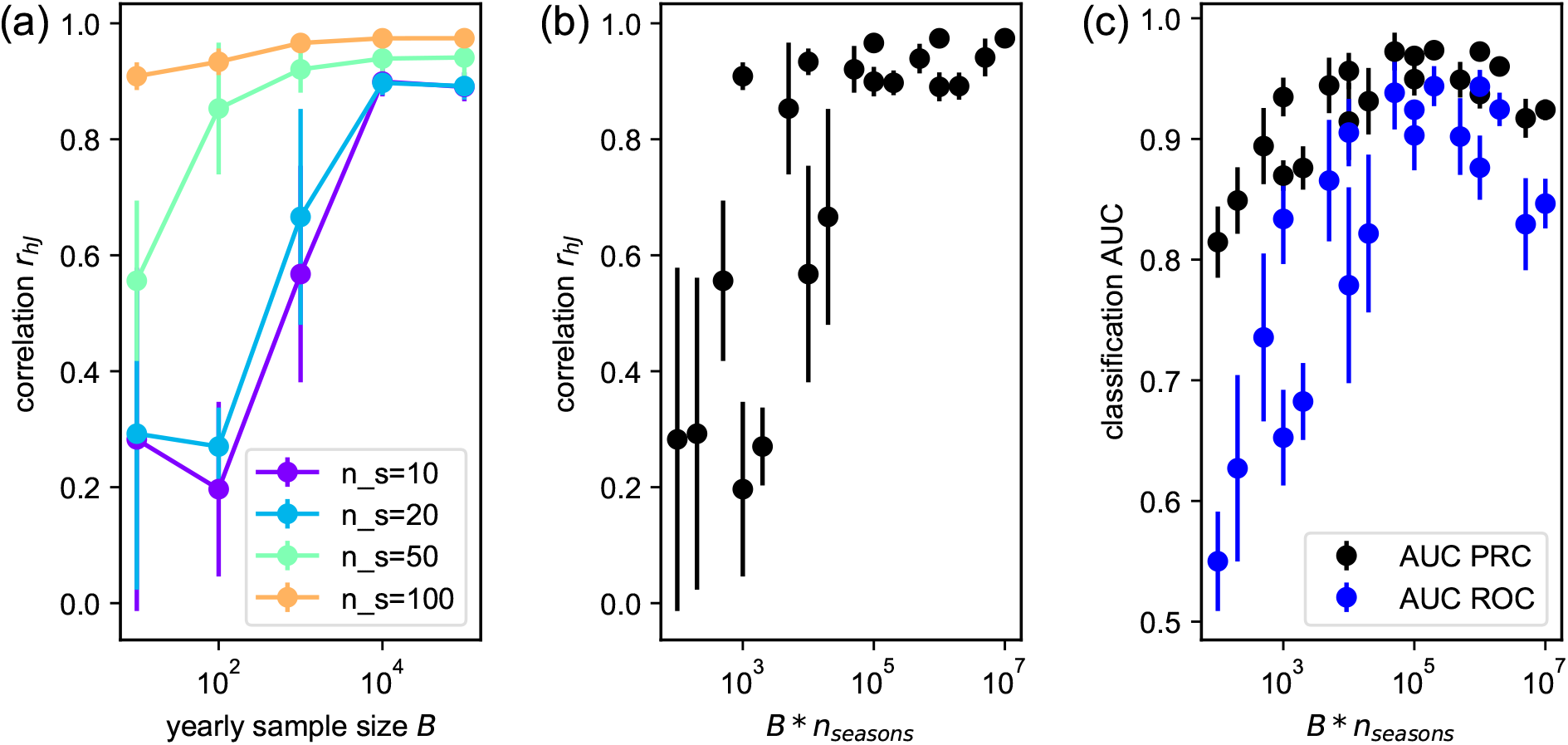
Inference performance for varying yearly sample size *B* per season and varying number *n*_seasons_ of seasons used for inference. (a) The correlation coefficient *r*_*hJ*_ between inferred and simulated double-mutational fitness costs as function of yearly sample size *B* for various *n*_seasons_. (b) The correlation coefficient *r*_*hJ*_ as function of total sample size *B* * *n*_seasons_. (b) The area (AUC) under the ROC curve and under the precision-recall curve (PRC) for classification of deleterious double mutations with classification threshold *F*_threshold_ = −10, shown as function of total sample size *B* * *n*_seasons_. Each value is averaged over 6 simulations, respectively, and errorbars show the respective sample standard deviations. For the shown example the fixed parameter values for simulation and analysis are: *N*_pop_ = 10^5^, *L* = 20, *µ* = 10^*−*4^, *σ*_h_ = 1, *D*_0_ = 5, *N*_simu_ = 200, *λ*_*h*_ = 10^*−*4^, *λ*_*J*_ = 1, *λ*_*F*_ *\** = 10^*−*4^.

Regarding the amount of data needed for fitness inference of a given antigen, we can quantify our rough scaling expectations with the following argument. For a sequence of length *L* and *n*_seasons_ the number of observed epidemic seasons, we need to determine *m* = *n*_seasons_ + *L*(*L* + 1)*/*2 ≈ *n*_seasons_ + *L*^2^*/*2 parameters. With the simplifying assumption that we need *m* independent equations for this inference task, we can estimate that we approach optimal inference performance when *B n*_seasons_ *µ* ∼ *m*. Here the number of needed total samples *B* * *n*_seasons_ is assumed to increase with decreasing mutation rate *µ*, since an independent set of samples is obtained only roughly every 1*/µ* years.

The inference performance further strongly depends on other parameters such as the sequence length *L* (Fig. 8a) and on the population size *N*_pop_ (Fig. 8b). Inference performance in terms of the correlation *r*_hJ_ between inferred and simulated double-mutational fitness coefficients decreases with increasing sequence length and increases with increasing population size towards an upper limit ≤ 1. Thus, if the protein sequence representation is high-dimensional, a very large amount of data is needed for a high inference performance (indicating the need for dimensionality reduction) and secondly, if the effective population size that defines the selection bottleneck is small, inference can be poor. A large population size, however, will not help to have high inference performance if the sample size *B* is low.

**Figure 8.**
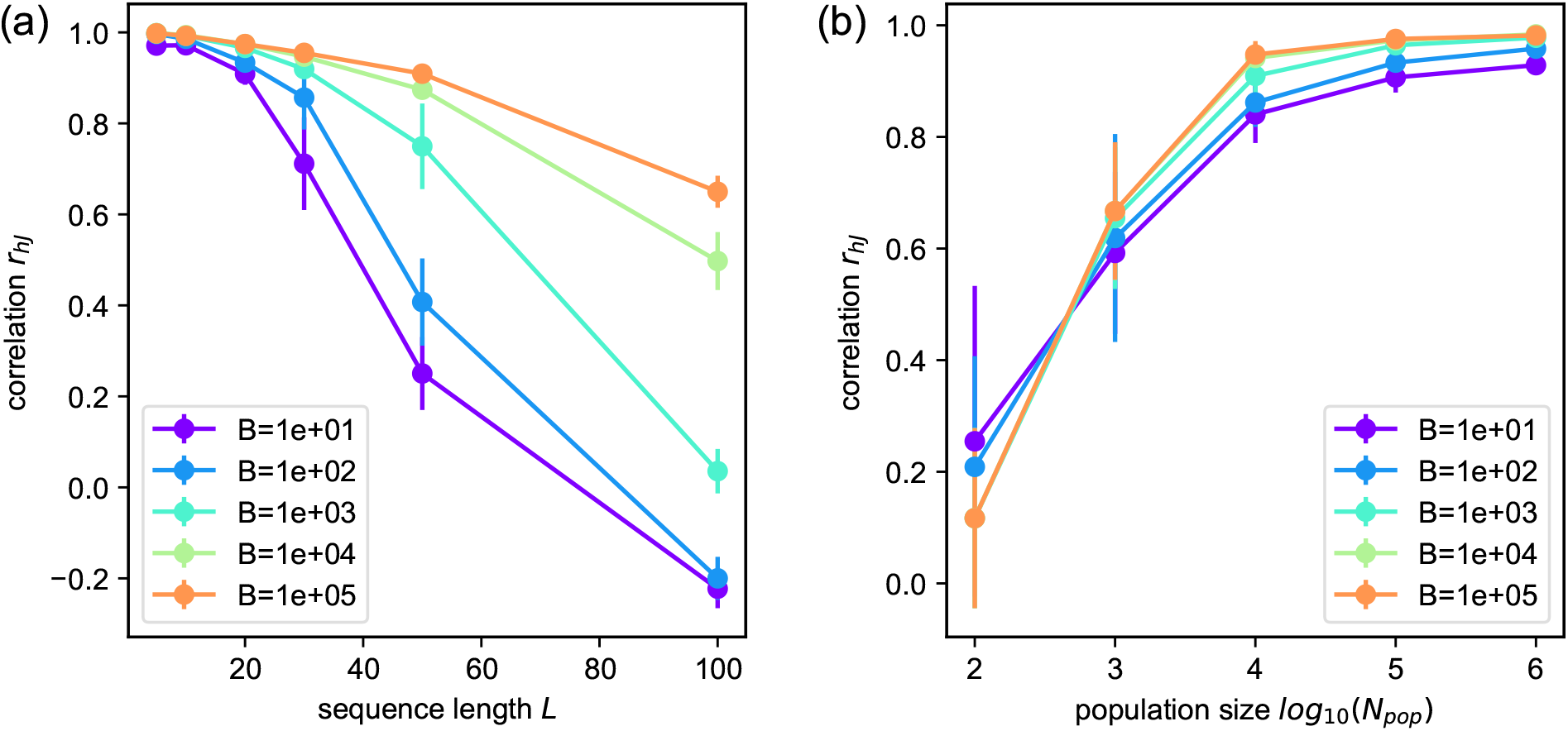
Inference performance in terms of the correlation coefficient *r*_*hJ*_ between inferred and simulated double-mutational fitness costs, for varying simulation and analysis parameters. (A) Inference performance as function of sequence length *L*, (B) inference performance as function of population size *N*_pop_. For both parameter explorations the yearly sample size *B* was varied between 10 and 10^5^. Each value is averaged over 6 simulations, respectively, and errorbars show the respective sample standard deviations. For the shown simulation results the respective fixed parameter values for simulation and analysis are: *N*_pop_ = 10^5^, *L* = 20, *µ* = 10^*−*4^, *σ*_h_ = 1, *D*_0_ = 5, *N*_simu_ = 200, *n*_seasons_ = 100, *λ*_*h*_ = 10^*−*4^, *λ*_*J*_ = 1, *λ*_*F*_ *\** = 10^*−*4^.

## IV. DISCUSSION

Here we presented a method for inferring the intrinsic mutational fitness landscape of influenza-like antigens from population-level protein sequence time series data. Our approach is able to infer single as well as pairwise mutational effects for binary sequences with several tens of sites. By simulating influenza-like evolutionary dynamics, we were able to analyze inference performance under different conditions such as for various sequence lengths and sample sizes. Our inference approach, in principle, only relies on the raw strain frequency data as function of time and does not depend on a separate inference of sequence phylogenies, in contrast to other analyses [10, 11].

In comparison to the recently proposed marginal path likelihood method (MPL) for sequence time series [54], we were able to disentangle time-varying immunity-dependent fitness effects from intrinsic fitness, and we not only inferred the fitness effects of single mutations but also of double mutations at pairs of sites.

In order to make meaningful predictions based on observed influenza protein sequence data, our inference approach needs to be translated to this more complex system, which generally has a high-dimensional sequence landscape with around hundred residues in the head epitope regions of HA (A/H3N2) and 20 possible amino acids per residue. The inference performance will also be constrained by a relatively small number of samples, around 3 * 10^4^ HA sequences in total between 1968 and 2020 [63, 64].

For using our inference approach on the influenza protein data, one further needs to make sure that the cross-immunity function in −*F*_host_ (Eq. 3), which we use as response variable, adequately captures the cross-immunity between different strains. The total mutational distance in the epitope regions, which we use in our model and which has been used in previous studies [10] for estimating cross-immunity, only roughly captures the cross-immunity measurements from hemagglutination inhibition (HI) assays [6, 65]. Analysis of such HI data, in which the proposed cross-immunity function is compared against measured cross-immunities, suggests a typical cross-immunity distance *D*_0_ of 5 amino acids or 14 nucleotide residues for seasonal influenza A (H3N2) strains [10, 65], i.e., two strains that differ by more than 5 amino acid mutations within their epitope regions typically experience negligible cross-immunity to each other’s immune responses.

For testing fitness inference performance on real data, we generally do not have much direct information on the intrinsic effects of various mutations besides from some in-vitro mutational assays, which are locally constrained to small parts of the sequence space or which only consider single-mutational fitness effects based on a given reference strain [66–69]. Furthermore, these empirical studies only measure fitness in terms of functional replication in cells, not in terms of spread across the human population. The application of classical machine-learning methods of testing inference based on predictions on held-out data are also challenging due to the complex time-dependent nature and general sparsity and heterogeneity of available sequence data.

Our computer simulations with a well-defined model of the evolution of a mutable virus subjected to human immune pressure over time have generated a data set of temporally ordered sequences. These sequences could be used in the future to test the veracity of different inference schemes against data that is the “ground truth”. For example, do existing models developed for predicting the most likely influenza strains given data until the preceding year [10, 11] give the right answers for the data set that we have generated?

In conclusion we have proposed a method for inferring the intrinsic mutational fitness landscape of influenza-like viruses from time series of observed antigenic sequences. This approach can, with increasing availability of sequence data in the future, contribute to the development of new cross-and long-term protective immunization strategies against viruses that evolve due to immune-driven selection. Like seasonal influenza, SARS-CoV-2 and other novel viruses might become endemic by evolving under vaccine and natural immune pressure, and our approach might provide valuable insight into their intrinsic fitness landscapes and reveal their vulnerabilities.

## CODE AND DATA DEPOSITION

Computer code used for simulations and analyses as well as data that were used to produce the figures in the paper are available on GitHub at https://github.com/JDoelger/InfluenzaFitnessInference.git.

## ACKNOWLEDGMENTS

This research was supported by the Ragon Institute of MGH, MIT, and Harvard and NIH grant # R01HL120724-01A1. MK acknowledges support by the NSF through grants # DMR-1708280 and # PHY-2026995.

## AUTHOR CONTRIBUTIONS

AKC and MK initiated the project. JD, MK and AKC designed research, analyzed the results and wrote the paper. JD performed the calculations and generated all figures.

The authors declare no competing interests.

